# Decline and extirpation of birds in a tropical Asian grassland ecosystem over the past century

**DOI:** 10.64898/2026.11.09.765392

**Authors:** Sutirtha Lahiri, Jugal Borah, Sushma Reddy

## Abstract

Grasslands are facing unprecedented habitat loss due to settler-driven agriculture during colonial expansion, conversion to tree plantations, misclassification as wastelands, absorption into megadevelopment projects, and insufficient response to climate change. Globally, grassland birds have faced the steepest declines, with many formerly common species facing rapid decline, losing much of their original ranges. In South Asia, our knowledge of how these land-use changes in grasslands have affected birds remain scant. Natural history collections and archives are vital resources to identify historical baselines for conducting a resurvey and identifying trends in long-term biodiversity response. Yet such historical data remain insufficient in tropical regions. In this study, by compiling an unprecedented dataset of ornithological surveys from the 1900s using a diverse set of historical data, we conducted a 3-year resurvey in an Eastern Himalayan grassland first surveyed by British Ornithologist Herbert Stevens. We surveyed nine historical localities. Using a recently developed Bayesian method to calibrate modern surveys with historical museum data, we found a 70.5% decline in grassland specialist birds. We also report the local extirpation of four endemic species of grassland birds. Open habitat raptors showed a 74.9% decline, while generalist species showed an increase of 23.5% and wetland birds showed a marginal increase of 2.64%. By quantifying functional distinctiveness, a measure of morphological diversity, we find that loss of grassland species does not impact morphological and functional diversity in modern assemblage. We also find that most resurvey localities have a high number of invasive species that have completely altered these communities. Grassland specialist birds of South Asia are highly range-restricted and have specific habitat requirements; the lack of intact tall grass communities in concert with site-level disturbance pressures from grazing, conversion and fire have likely contributed to their local extirpation. Our study serves as one of the first efforts to resurvey wet grasslands of South Asia and document local-level extirpation. We also highlight the importance of museum-based research to document long-term changes for urgent conservation attention.

## Introduction

Grasslands are one of the most at-risk habitats, facing rampant loss due to anthropogenic impacts (Rosenberg et al. 2019; Douglas et al. 2023). In South Asia, there has been a loss of an estimated 20 million hectares (45%) of grassland between 1880 and 2010 (Tian et al. 2014), leading to severely fragmented patches of this habitat. Yet many of these grasslands continue to remain poorly studied because they are not as species-rich as forests. Conserving landscapes such as grasslands requires understanding patterns of temporal change in diversity and extinction, which can be possible by historical baselines. This study ties conservation priorities of birds and historical ecology to ask: *How has historic loss of the grasslands affected patterns of bird diversity and extinction*?

Grasslands worldwide have been lost due to unprecedented conversion during European colonization (Murphy et al. 2016). The European settlers saw immense potential in the vast, nutrient rich lands and sought to convert these habitats into productive landscapes. The expansion of agriculture, followed by industries and railroads, collectively led to a steep decline in old-growth grasslands. In North America, more than 99% of prairie grasslands were converted to agriculture, with similar trends in other major grasslands like the Cerrado (Silveira et al. 2022) in South America, the savannas of Africa and the semi-arid and wet grasslands of Asia (Ratnam et al. 2016a; Joshi et al. 2018). In addition to agriculture, large scale expansion of tree plantations (Roy et al. 2015; Banerjee et al. 2023; Mhetre et al. 2026), invasive species (Sinha et al. 2022) unscientific management (Kher & Dutta 2021), mining, megadevelopment projects and climate change impacts (Joyce et al. 2016) have collectively threatened grassland ecosystems. Additionally, misclassification of grasslands as degraded forests or ‘wastelands’ have further led to their decline (Lahiri et al. 2023).

The Himalayan foothills grasslands (hereafter wet grasslands) have undergone precipitous fragmentation, and the effect on birds of the region is acutely understudied. Rich in biodiversity and endemism (Lahiri et al. 2021a), the foothills formerly possessed extensive tall alluvial grasslands maintained by flood regimes of the Brahmaputra, Ganga and Indus Rivers and its tributaries. Wet grasslands have declined considerably in extent and quality due to large-scale conversion to tea plantations, agricultural lands during the green revolution as well as habitat loss due to the changing hydrology of rivers. These grasslands currently occur as fragmented habitats in a matrix of multiple land use (Rawat & Adhikari 2015; Banerjee et al. 2023). The wet grasslands remain ornithologically poorly explored despite harboring endemic bird species (Rahmani et al. 2016). The impact of historical and ongoing grassland habitat loss on endemic bird species—many of which are strict grassland specialists and of high conservation concern (Baral 2001; Rahmani et al. 2016; Warudkar et al. 2022a; Jahan et al. 2022, 2023; Sharma et al. 2026) —require urgent study as they face continued threats due to climate change, small population sizes and human exploitation (Lahiri & Reddy 2025).

Globally, grassland birds face the steepest declines (Rosenberg et al. 2019; Ogada et al. 2022; Shaw et al. 2024) compared to other biomes. In the Indian subcontinent, several grassland species are threatened and likely locally extirpated from a large part of their historical range (Ganesh & Prashanth 2018; Kher & Dutta 2021; Bharadwaj et al. 2024; Ramesh et al. 2025; Mhetre et al. 2026). In the wet grass grasslands, too, birds have faced considerable decline (Jha et al. 2018; Lahiri et al. 2021a; Jahan et al. 2022; Choudhury 2023). Wet grasslands consist of short and tall grasslands. While short grasslands (primarily *Imperata sp*.) occur in drier soil and require fire management and often occur, tall grasslands (*Phragmites- Arundo- Saccharum*) occur in wetter, sandy soil with frequent flooding (Lahiri et al. 2021a; Jahan et al. 2022). Wet grassland birds have very specific microhabitat requirements, further aggravating their propensity to decline. For example, while species like Marsh Babbler *Pellorneum palustre* require tall stands of inundated *Phragmites-Arundo* grasslands, Indian Grassbird *Graminicola bengalensis* and Bengal Florican *Houbaropsis bengalensis* require shorter *Imperata* grasslands. The Black-breasted Parrotbill (*Paradoxornis flavirostris)* has been extirpated from much of its former range, and exists in only a few scattered, and likely unconnected, localities (Krishnan 2021). Similarly, the Bengal Florican *Houbaropsis bengalensis*, a species of shorter wet grassland, is critically endangered and number fewer than 300 in India.

Natural history museums have long documented biodiversity changes over time and space. More recently, studies have demonstrated their relevance for conservation by using museum specimen records to show biodiversity loss (Norris et al. 2024). By generating historical baselines from museum expeditions, studies have used a historical resurvey framework to analyze changes in species richness and community composition (Palacio et al. 2020; Choi et al. 2020), abundance (Ramesh et al. 2025), loss of genetic diversity (Benham et al. 2024), changes in migration phenology (Gutiérrez-Carrillo et al. 2024) and local extinctions (Cavarzere et al. 2017; Gómez et al. 2021). However, while long-term, and replicable, historical datasets exist in developed countries and have allowed for robust estimation of species trajectories (Magurran et al. 2010; Iknayan & Beissinger 2018), such data are severely lacking in tropical regions where data were collected primarily for opportunistic species documentation, discovery or sport while collectors were posted in administrative services (Guneratne 2023). Expeditions in tropical regions were also often impeded by complex geopolitical pressures, including war and lack of access to permits. As a result, replicable historical data are lacking, and collating such data from multiple sources is imperative to quantify biodiversity changes across longer time periods.

In this study, we combine historical records from natural history museums with a 3-year systematic bird survey to ask-a) How have species relative abundances shifted across a century? b) Are species that have been lost/extirpated morphologically and functionally unique? We surveyed a total of 54 point count stations in 9 historical localities in the Subansiri floodplains of Assam, Eastern India and quantified and compared species relative abundance across the two time periods. By undertaking this study which is, to the best of our knowledge, the first resurvey effort focused exclusively on grasslands in the Himalayan region, we aimed to quantify the extent of species population changes across a century of habitat loss. Our study also aimed to demonstrate the utility of museum-based resurveys that are still underrepresented in tropical countries owing to limited historical data and reduced access to museums and archives (Ocampo-Peñuela et al. 2026).

## Methods

### Historical survey

Due to lack of standardized data, we first conducted an extensive archival survey to compile a historical dataset from multiple sources. One of the key considerations of an acceptable historical dataset is that it should have species, date and location information available. Because we did not know of any expeditions specifically conducted in grasslands, we first made a list of grassland specialist birds from South Asia (following the eBird/Clements taxonomy) that are of conservation concern, and searched museum databases such as Vertnet, iDigBio as well as museum-specific databases. The following museums had substantial collections: Bombay Natural History Society (BNHS), American Museum of Natural History (AMNH), The Great North Museum, Newcastle Upon Tyne, Yale Peabody Museum (YPM), National Museum of Natural History (USNM), Museum of Comparative Zoology, Harvard (MCZ), Royal Ontario Museum (ROM), Museum für Naturkunde Berlin, Natural History Museum, Tring. For museums that did not have an online repository, such as Natural History Museum, UK and Bombay Natural History Society, we visited the museum collections to collect data. For each specimen, we recorded several attributes such as location, year of capture, collector and any other additional information present on the tags. From this metadata, we then traced additional species that were collected on the same expeditions. For example, once we had a set of names of collectors, we queried archives and libraries for archival data, such as journals, letters, correspondence, and any other form of historical material. We perused these archival materials to look for any species checklists, locality information, description of sites, maps and other information.

From this triangulation method, we finally selected the ornithological expedition of Herbert Stevens as a good representation of baseline surveys. Herbert Stevens was a British tea planter who spent a decade (1900-1910) in the Lakhimpur district of Assam. His dataset contained an extensive list of species, which includes detailed notes on locations, dates as well as numbers. In addition to specimens collected by him, he also made accounts of species he saw during the course of his stay in the district. While ornithological expeditions are usually biased towards the collection of rarer specimens (the rarity-seeking syndrome) (Reddy and Davalos 2003), Stevens made detailed observations and collections of both rare and common species. Thus, his expedition was more of an extensive avian diversity assessment than selective collection of rare birds. We digitized his specimen records from museums such as the Newcastle Museum, Natural History Museum (UK), Bombay Natural History Society, American Museum of Natural History, Yale Peabody Museum. We also digitized his observational records from his journal article (Stevens 1915). Since Lakhimpur district encompassed a wider region in the 1900, we limited our survey to only include present day Lakhimpur district.

### Study Area

Lakhimpur district is situated in Upper Assam, bordered by the Himalayan range in the north, and flooded by the River Subansiri, one of the main tributaries of the Brahmaputra River (Fig. 2). Other smaller rivers include the Dikrong and the Ranganadi. The floodplain is situated at an elevation of 200 msl and consists of diverse land-use including floodplain grasslands, tropical deciduous forests, wetlands, tea plantations, agriculture, and grazing commons. Our study sites, specifically, were in the grasslands on the riverbanks of the Subansiri, Dikrong and Ranganadi rivers.

**Fig 1:**
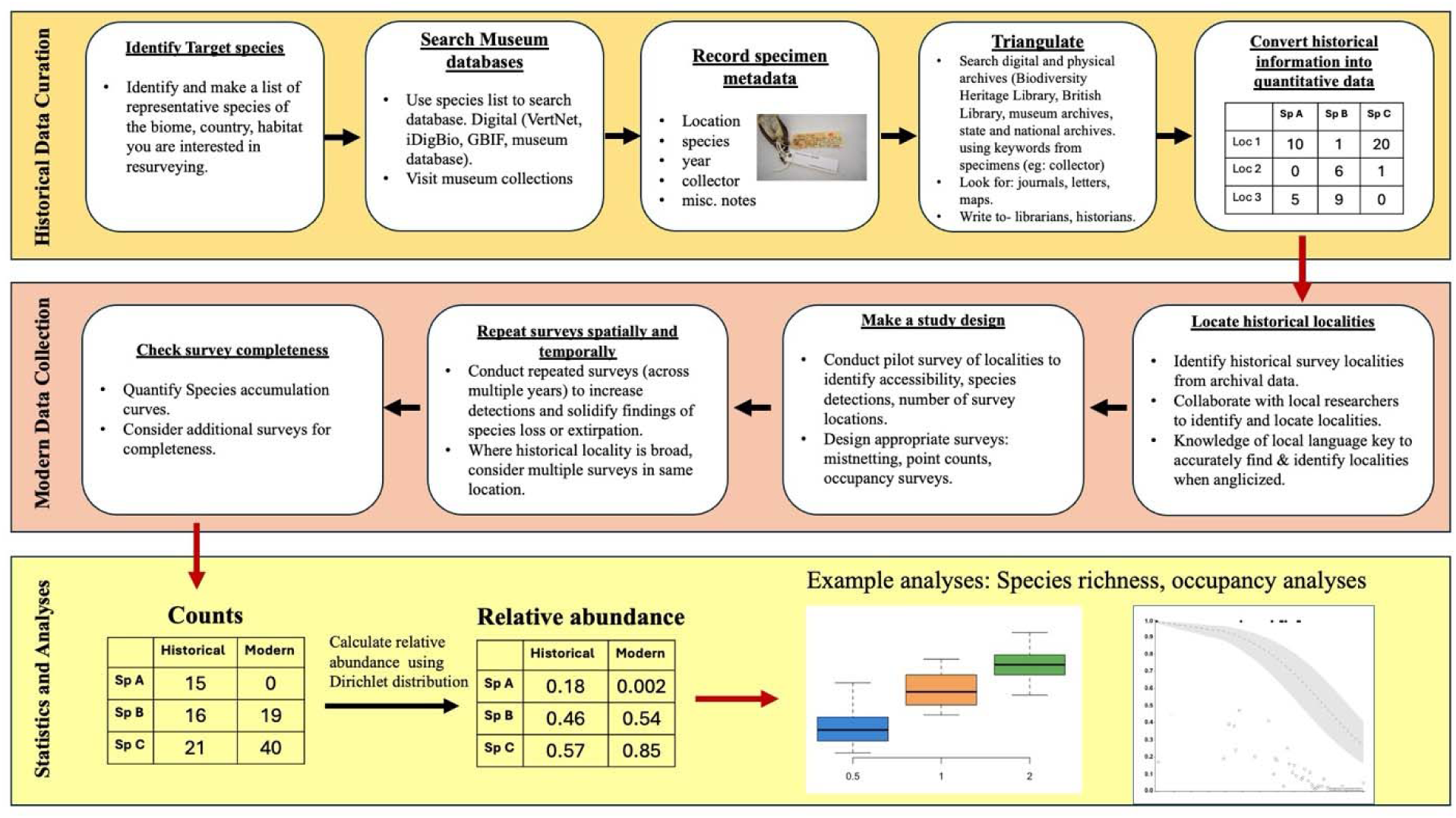
A replicable workflow to identify, curate and collect historical datasets from NHMs, archives and digital repositories to conduct a resurvey.

**Fig 2:**
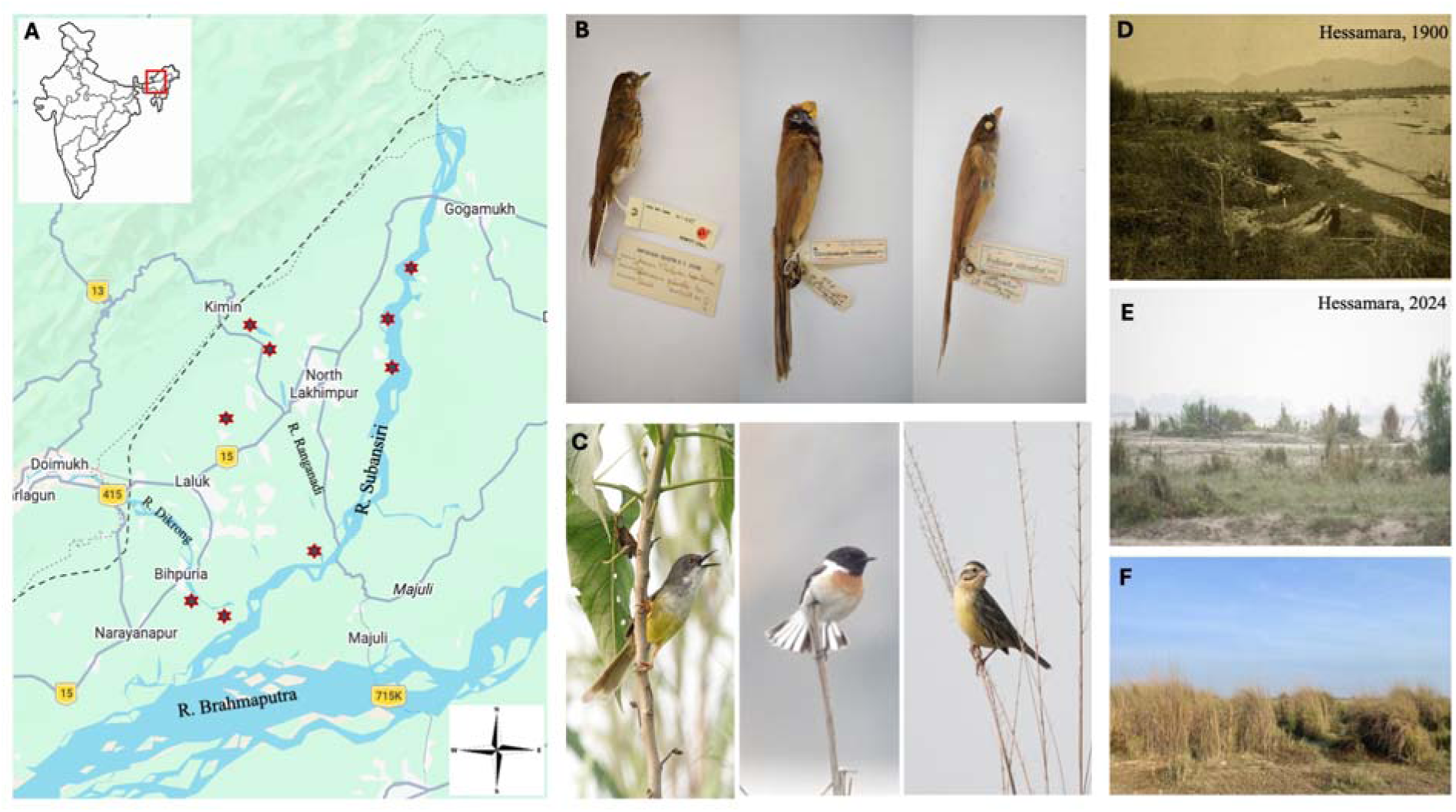
A. Map of study area. The study was conducted at 9 sites along the Subansiri, Dikrong and Ranganadi rivers. The red stars denote our study sites. B. grassland specialist birds (from L-R) *Pellorneum palustre* (VU), *Paradoxornis flavirostris* (VU) and *Chrysomma altirostre* (VU) recorded by Herbert Stevens C. Grassland specialist birds *Prinia flaviventris, Saxicola maurus*, *Emberiza aureola* recorded in our survey D. Historical photo of Hessamara, with tall grass species such as *Saccharum ravennae, Arundo donax, Phragmites karka* E. Modern photo of Hessamara. Grassland dominated by *Saccharum spontaneum* and *Tamarix dioica* in a matrix with agricultural fields and heavy sand deposits. F. Boduti grassland with stands of *Saccharum spontaneum* grass. Map lines delineate study areas and do not necessarily depict accepted national boundaries.

### Field survey

We conducted detailed bird surveys across three winter sessions (2024-2026) and one breeding season (2024) in sites which were surveyed by Stevens using variable-distance point counts. At each of our 9 localities, we had 6 point count stations, for a total of 54 point count stations. We were unable to keep a consistent number of point count stations at each site owing to the small size of the grasslands, and lack of accessibility. In some sites, we had to shift our point count locations because our earlier locations were washed off during the monsoon.

Each point count station was spaced 150m apart. We conducted counts for 15 minutes, recording every species seen or verifiably heard. We also included species that flew over, as we were interested in detecting all species. In addition to bird counts, we also took habitat and site variables, such as grass height, distance from water, presence of invasive species, shrubs as well as anthropogenic activities such as fire, grazing and cutting. For the anthropogenic activities, we coded them from 0-2, with 0 being no activity, 1 for intermediate, and 2 for severe. These coding were vetted by both SL and JB to remove individual bias. At each point count location, we also took measurements of grass height and dominant grass species.

To assess survey completeness, we calculated metrics of species diversity as well as species accumulation curves using the package vegan (Oksanen et al. 2018) and iNext (Hsieh et al. 2016) in R programming language (R core team 2025).

### Statistical analysis

#### Relative abundance

We first coded each species as belonging to one habitat category. We did so by first using the habitat categories from AVONET (Tobias et al. 2022) and recoding each species into one of four categories; grassland species, generalist, open habitat raptor and river/sand/wetland specialist. To re-code habitat, we relied on existing natural history literature (Handbook of the Birds of the World, (Ali & Ripley 1968; Grimmett et al. 1999; Rahmani et al. 2016) as well as field observations. We believe recoding was necessary as many of the available habitat categories for the birds in the study area were coarse scale.

We filtered all historical species with less than 5 occurrences. This filtering pruned rare species, thus retaining a total of 42 species of birds from a total of 124 species. For our field surveys, we truncated all species detected >100m away. Next, to calculate species relative abundances, we used the FAMA (Field abundance- Museum abundances) method (Gotelli et al. 2023). The method fits species count data (both historical and modern) into a Dirichlet distribution to ensure that the probability of occurrence for all species was >0, even if it wasn’t detected at any time period. The Dirichlet distribution uses a Bayesian method to handle zeros, whereby species that are not detected at any time period have an infinitely small non-zero value, whereas frequentist relative abundance for a non-detected species is zero, which is often not the case (Gotelli et al. 2023). We used the *rdirichlet* function in the *gtools* library (Lumley & Warnes 2010) to simulate 10000 relative abundance values for each species, as well as 95% CI.

Historical surveys have often focused on rarer species (Gotelli et al. 2023), thereby inflating the number of specimens. We checked if including rarer species might potentially skew the calculation of historical abundance. To do so, we re-ran our analyses by removing 4 endemic grassland species we believe could possibly skew our data. These include *Paradoxornis flavirostris, Pellorneum palustre, Laticilla cinerascens, Chrysomma altirostre*. In addition, we also removed one generalist species that had an exceptionally high modern abundance, *Saxicola maurus*. We did so to ensure that the relative abundance calculations were not skewed by one hyper abundant species. We used Welsh’s t-test to compare if species relative abundances differed between historical and modern time periods.

We calculated log response ratios (LRR) for each species in order to quantify the magnitude of change in relative abundance in each species. We defined log response ratio as the log (historical relative abundance/modern relative abundance). We calculated the credible intervals of the LRR by taking the logarithmic upper and lower CI outputs from our FAMA analyses.

#### Trait-based analyses

To explore how functional diversity has changed over time, we compiled morphological traits from AVONET (Tobias et al. 2022), including beak length, beak width, tarsus length, tail length, wing length, hand-wing index and mass. We standardized the data by first log-transforming mass, and then Z-transforming all of the traits such that the dataset has a mean of 0 and a standard deviation of 1 following established methodology (Gomez et al. 2021). Following this, we built a matrix containing species that have been extirpated and species that are shared between the two time periods. We also combined measures of relative abundance into this data frame. We calculated functional distinctiveness using a gower distance-based distance matrix (Gomez et al. 2021). We then calculated functional distinctiveness using the ‘distinctiveness_stack’ function in R package funrar (Grenié et al. 2017), weighted by relative abundance. We coded the each species as ‘shared’, if they are present in both historical and modern samples, ‘naive absence’ if they were historically present but absent from our surveys, and ‘locally extirpated’, if they were historically present but absent from both our surveys as well as other modern surveys and birdwatching reports.

## Results

We conducted 193 point counts in 2024, 2025 and 2026, recording 4708 individuals of 177 species. Species richness across each year was 130, 114 and 95 species, respectively (Table S1, Fig S1). Our sampling effort documented bird species with sufficient efficiency. Species accumulation curves show a sampling completeness of 99% across all our sites (Fig S1). *Tamarix dioica* was present in 33% of our sites, while the predominant tall grass species in all our sites was *Saccharum spontaneum* with a mean grass height of 0.55 ± 0.11m. 78% sites had evidence of cutting, 100% had evidence of grazing and 44% had evidence of fire (Fig S2).

Overall relative abundance of all species was significantly reduced between time periods (p<0.005) [Table 1, Figure 3]. Grassland specialist birds show a 70.5% decline, with 17 out of 20 (85%) grassland specialists showing a decline [Fig 4]. Among the species with the steepest declines, four species of grassland specialist birds that are endemic to South Asia-*Paradoxornis flavirostris, Pellorneum palustre, Laticilla cinerascens, Chrysomma altirostre-* have been locally extirpated. The decline in grassland specialist birds remains unchanged even when we remove these four species of birds. Open habitat raptors show a 74.9% decline (p<0.05) [table 1].

**Fig 3.**
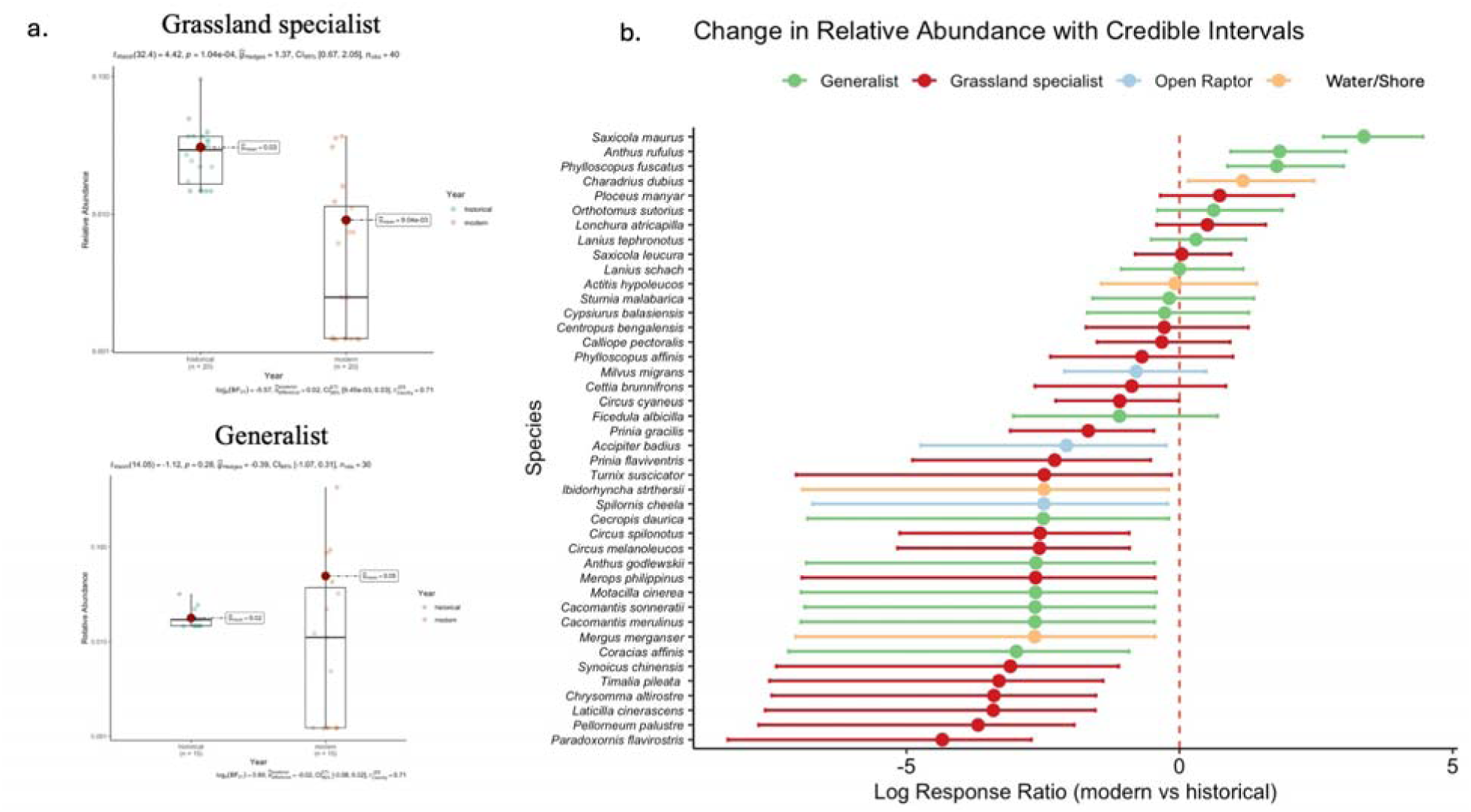
Panel a) boxplots of species relative abundances for grassland specialist and generalist birds. Grassland specialists show significant reduction in overall relative abundance compared to historical time. Figure b) log response ratio of each species along with their credible intervals calculated as an output from the FAMA analyses. The dashed line at 0 represents no net change in log response ratio, with overlapping bars representing non-significant change, and non-overlapping bars representing significant changes to LRR in species.

**Fig 4:**
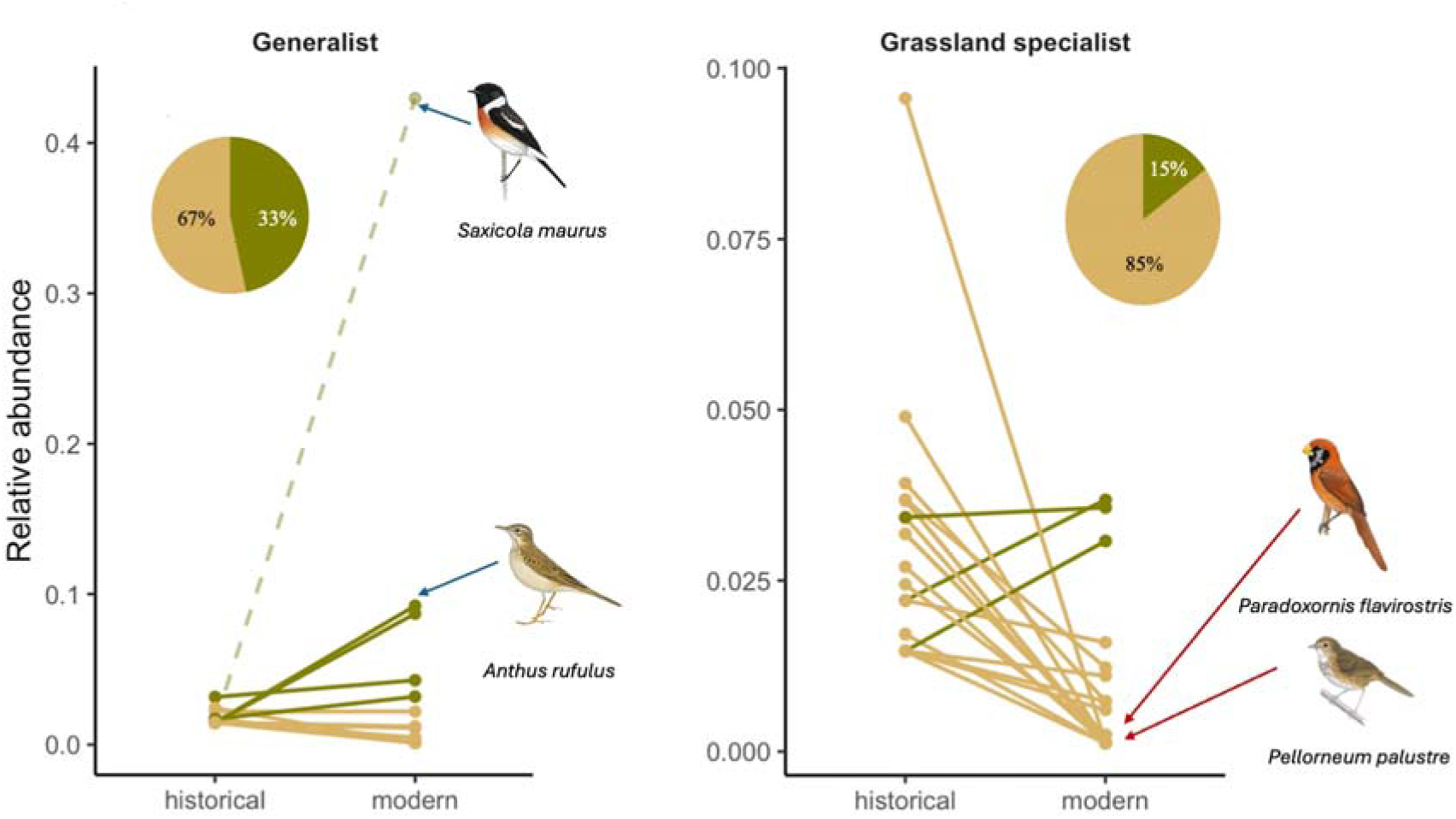
Mean relative abundance for generalist and grassland specialist species is shown. Lines in green represent an increase while lines in khaki indicate decline. Pie charts are the proportion of species of each habitat category that have declined or increased. The increasing trajectory of *Saxicola maurus* is shown using dotted lines since its mean abundance is much higher than the other generalist species. Illustrations used with permission from birdsoftheworld.com.

**Table 1:** Change in relative abundance of birds in each habitat category. % change (all species) refers to changes in relative abundance when we do not exclude the 4 grassland and 1 generalist species from our dataset, while % change (filtered) is the change when we exclude those species from our analyses. Filtering rarer species from our analyses does not change overall trends in the habitat categories. The font in red denotes significant changes.

| **Habitat** | **% change (all sp)** | **% change (filtered)** | **Mean LRR** | **CI low** | **CI high** |
|---|---|---|---|---|---|
| **Grassland specialist** | -70.50 | -55.4 | -1.87 | -2.58 | -1.16 |
| **Generalist** | 177 | 23.5 | -0.646 | -1.76 | 0.466 |
| **Open Raptor** | -74.9 | -74.9 | -1.78 | -3.97 | 0.401 |
| **River/sand/wetland** | 2.64 | 2.64 | -1.01 | -3.98 | 1.96 |

Generalist species showed an increase of 177% in total, despite 10 out of 15 generalist species showing a decline [Fig 3]. The species with the highest increase is *Saxicola maurus,* followed by *Anthus rufulus* and *Phylloscopus fuscatus*. Since one species, the *Saxicola maurus,* disproportionately skewed our results, we also re-ran our analyses by removing this species. We found that removing *Saxicola maurus* from our dataset showed a modest, but statistically insignificant, increase of 23.5% in generalist birds (Table 1). Wetland birds showed a marginal, but statistically insignificant, increase of 2.64% (p>0.05). Our log response ratio also corroborated our relative abundance results, showing statistically significant declines in grassland specialist birds (LRR –1.87; CI −2.58 – −1.16, table 1).

Functional distinctiveness analyses showed no significant difference between naive absence and shared species assemblages (p>0.05). Within the naive absence, there were four species of grassland specialist birds- *Paradoxornis flavirostris, Pellorneum palustre, Laticilla cinerascens, Chrysomma altirostre-* and one species of riverine bird- *Ibidorhyncha struthersii*- that we categorise as ‘locally extirpated’. The species that have been locally extirpated are not functionally distinct but are rather situated within the functional space of other species in the assemblage [Fig 5].

**Fig 5:**
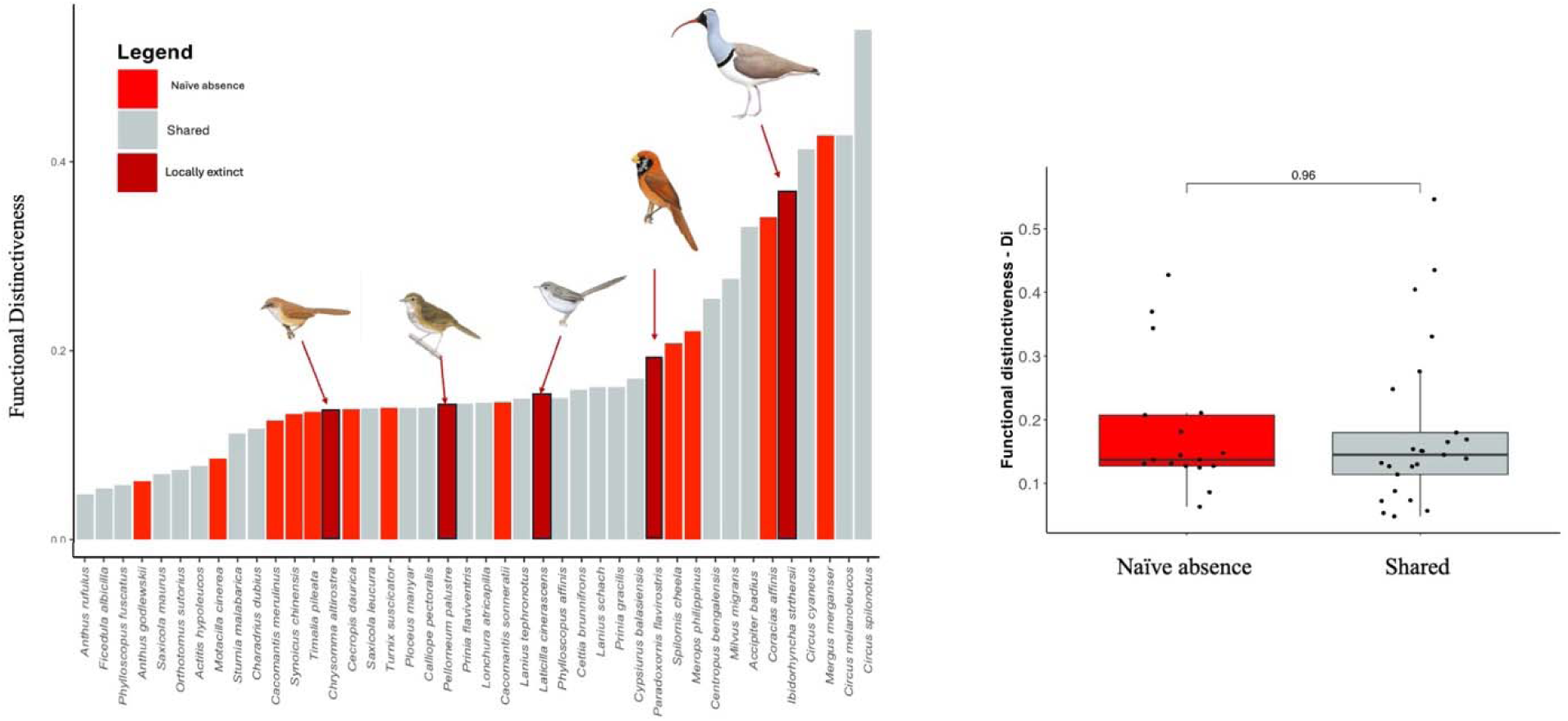
Functional distinctiveness of naive extinctions (red) and shared species (gray) for the Subansiri Floodplains, India. Difference between the two categories is statistically insignificant (p=0.96). The locally extinct grassland species (represented by dark red bars- *Paradoxornis flavirostris, Pellorneum palustre, Chrysomma altirostre, Laticilla cinerascens*) are also not functionally distinct, being medium- sized birds. The only extirpated species that has higher functionally distinctiveness is the Ibisbill, a riverine species. Species illustrations used with permission from birdsoftheworld.com.

## Discussion

To summarize, using a historical resurvey approach, we report the decline in species across major habitat guilds. Grassland specialist birds show the most significant decline of 70.5%, while open habitat raptors decline by almost 75%. Concomitantly, we report the rise in abundance of generalist species like *Saxicola maurus, Anthus rufulus* and *Phylloscopus fuscatus*. Our historical dataset, gathered from museums and archives, provide a valuable benchmark for comparing and substantiating context for current biodiversity loss.

Grassland birds have faced the highest declines globally (Veldman et al. 2019). In the high- altitude shola grasslands of South India, studies have reported declines in grassland specialist birds (Ramesh et al. 2025) due to conversion to agriculture and timber plantation. Our findings are consistent with global trends, with a 70% decline of grassland specialist birds, and a 75% decline in open habitat raptors in the Subansiri floodplains. In addition to the decline and extirpation in endemic species, we also find widespread decline in otherwise common grassland specialists and raptors like *Prinia flaviventris, Timalia pileata, Merops philippinus, Synoicus chinensis, Prinia gracilis, Circus cyaneus*, which were detected in our surveys, albeit in very low abundance. Open habitat raptors have shown decline population trajectories globally (Ganesh & Prashanth 2018; Shaw et al. 2024). Many of these declines have been linked to changing land-uses in both breeding and non-breeding sites, as well as the increasing use of pesticides in agricultural lands that lead to reduction in prey as well as mortality (Newton 2017).

Our finding of the local extirpation of the four species of grassland specialist birds raises alarm about the fate of the Himalayan lowland grassland biodiversity. Birds like the *Paradoxornis flavirostris, Pellorneum palustre, Laticilla cinerascens, Chrysomma altirostre* are highly range-restricted and occur in only wet grassland habitats within tall, dense and unbroken stands of *Phragmites, Arundo donax, Saccharum ravennae* and *Vetiveria sp.* grasslands (Krishnan 2021; Lahiri et al. 2021a; Warudkar et al. 2022b). These species have declined from much of their historical ranges. The Black-breasted Parrotbill, for example, has been locally extirpated from the West Bengal Dooars region, where they were formerly common residents (Krishnan 2021). Similarly, *Pellorneum palustre* and *Chrysomma altirostre* has declined from much of their former ranges and today occurs in only isolated grassland patches in the Northeast of India (Choudhury 2023). Local extirpation of grassland birds is evident across much of South and Southeast Asia. In the Western Ghats, the grassland specialist Nilgiri Pipit *Anthus nilghiriensis* has been locally extirpated from much of its former range (Lele et al. 2020). Grassland specialists like the great Indian Bustard, Lesser Florican, Bengal Florican, Broad-tailed Grassbird, Bristled Grassland have shown similar declines across their ranges (Dutta et al. 2011; Packman et al. 2014; Jha et al. 2018; Mahood et al. 2020; Kher & Dutta 2021; Bharadwaj et al. 2024). With local extinction and range reduction, grassland specialist birds are being pushed to isolated populations with concerns for inbreeding and reduced genetic diversity(Srinivas et al. 2022).

We also report the local extirpation of a riverine specialist species, the Ibisbill (*Ibidorhyncha struthersii).* The Ibisbill breeds in the high-altitude shingle-bed mountain river valleys of the Himalayan mountain range (Sinha et al. 2026) and comes to lower altitude, fast-flowing rivers during the winter. While not of conservation concern, they are tied to fast-flowing rivers during winters and are prone to disturbances like stone and gravel mining and hydropower dam-related disturbances.

Several studies have identified defaunation of large-bodied birds in response to historical land-use change (Gómez et al. 2021), however, our study highlights the loss of small-bodied birds. Ecosystems like flooded grasslands have a high diversity of species that are morphologically similar, as shown in our study. In conservation decision-making large-bodied, charismatic and colorful species are often accorded more focus in terms of research and funding. The lack of recognition of smaller-bodied species has, no doubt, led to a paucity of data on population trends and conservation assessments of many species, leading to local-level population loss and extirpation, such as the ones we have documented. The species of grassland specialists that have been locally extirpated are also ones with inconclusive and insufficient data for much of their life history. For many decades, these birds went undetected due to lack of any survey efforts. Today, we know very little about their nesting, movement, diet, habitat specificity, and responses to disturbance regimes. Thus, there is an urgent need for long-term and consistent monitoring efforts, including funding studies on fundamental biology of these species in order to aid in conservation decision making.

One of the reasons for the decline and local extirpation of grassland specialist birds is the loss of intact tallgrass habitat. Intact grasslands are crucial for the persistence of grassland specialists (Baral 2001; Rahmani et al. 2016; Warudkar et al. 2022a; Jahan et al. 2022, 2023; Sharma et al. 2026). Tall grasslands often exist as mosaics of drier patches of *Imperata* grasslands and wetter patches of *Phragmites-Arundo*. These mosaics present the required microhabitat that each grassland endemic depends on. *Paradoxornis flavirostris, Pellorneum palustre and Chrysomma altirostre* are exclusively tied to *Phragmites karka* and *Arundo donax* species that are tall and dense. In our study sites across the Lakhimpur landscape, we documented the near complete absence of the tallgrass habitats. Grasslands in the Lakhimpur landscape are smaller in area, occur in a matrix of agriculture and are dominated by *Saccharum spontaneum* and *Tamarix dioica*. Furthermore, fire, grazing and cutting are prevalent across all sites that we have surveyed. There is evidence of reduced nesting success, higher predation and declining abundance among wet grassland birds in other parts of Asia and Europe (e.g., *Panurus biarmicus*, *Paradoxornis heudei*)(Boulord et al. 2012; Barrionuevo et al. 2016; Ma et al. 2019). While very little is known about the effects of disturbance regimes on wet grasslands and biodiversity in India, studies have reported mixed responses. While some grassland birds persist and breed in habitats with moderate to minimal disturbance regimes such as grazing, cutting and fire (Baral 1999), other species do not prefer disturbance or unscientific management regimes such as untimely burning (Lahiri et al. 2021b).

Grasslands in the Lakhimpur landscape have been lost due to a variety of factors. While we did not test for the causality, our surveys across three years indicated that conversion to agriculture, erosion and siltation due to the changed hydrology of the rivers and the spread of *Tamarix dioica* have collectively posed the highest threats to these unique grasslands. Interviews with local community members and researchers also confirm that vast stretches of tall grasslands that were common in the landscape have been completely wiped out (Lahiri et al. in prep). Studies from other wet grassland sites show that changes in the hydrology of the river-due either to mega dam development or seismic activities- and the resulting siltation and erosion lead to changes in grassland establishment. While tall grass species like *Phragmites, Arundo* and *Saccharum ravennae* require moist soil and standing water, changing sedimentation promotes the expansion of *Saccharum spontaneum* and *Tamarix dioica*.

The industrial conversion of grasslands to agriculture and other land-use due to settler colonialism has been shown to be the most pervasive threats to these unique ecosystems (Ramesh et al. 2025, Vickery and Herkert 2001, Kher and Dutta 2019, 2026). Agriculture is one of the main occupations in nutrient-rich Himalayan floodplains. The Green Revolution of the 1960s further encouraged the conversion of wet grasslands to land suitable for mechanized agriculture of paddy and wheat (Strahorn 2009). Even though agricultural land has been shown to be important for grassland species, they cannot replace the value of intact grassland habitats (Kher and Dutta 2019). Nearly all of our grassland sites existed within a mosaic of agricultural land, many of which were actively being converted while our surveys were ongoing.

In addition to agriculture, the changing hydrology of the Subansiri and its tributaries due to large- scale hydropower dams upstream warrant urgent attention. Wet grasslands are maintained by the hydrology of rivers, as flooding and siltation ensure grassland regeneration as well as the control of woody encroachment and invasive species (Banerjee et al. 2023). Dams have already led to the decrease in grasslands, increased siltation and erosion in several parts of Northeast India, including in North Lakhimpur. While our study is focused on the floodplains of the Subansiri river, a similar trajectory of decline and local extinctions is evident in other wet grassland birds across their range. With countries focused on building the highest density of large-scale hydropower dams on Himalayan rivers (Grumbine & Pandit 2013), there is immense concern about their environmental and social impacts downstream, including the most imminent impact on floodplain grasslands that today house several endemic and at-risk species (Vagholikar & Das 2010; Sheth et al. 2020).

Our study also detected the presence and persistence of four species of bunting, including the critically endangered, Yellow-breasted Bunting *Emberiza aureola*. Buntings, despite being common, are facing severe population decline (Choi et al. 2023) across their range in Asia and Europe. Historical data has shown a marked change in bunting species composition across South Korea, with common species such as Meadow and Yellow-throated Buntings showing a yearly declining trajectory. Similar trends hold for many bunting species across their ranges in Europe and Central Asia, leading to 7 out of 44 species of buntings being listed as conservation concern. While the Yellow-breasted Bunting has faced a precipitous decline of 84.3 – 94.7% (Kamp et al. 2015), there is insufficient evidence for the population trends of the other three species of buntings that we have detected in our surveys. Being granivores, they depend on agricultural fields, in addition to grasslands, for foraging.

Additionally, we also detected multiple other grassland species of interest, including Swamp Francolin *Ortygornis gularis*, Chestnut-capped Babbler *Timalia pileata*, Striated Babbler *Argya earlei*, Jungle Babbler *Argya striata*, White-tailed Stonechat *Saxicola leucura*, Yellow-bellied Prinia *Prinia flaviventris*, Golden-headed Cisticola *Cisticola exodus*, West Himalayan Bush Warbler *Locustella kashmirensis*. While our study sites have suffered from massive land-use change that has led to local extirpation of grassland endemic species, they act as important stopover, wintering and breeding grounds for species like the *Ortygornis gularis*, *Timalia pileata, Argya earlei, Saxicola leucura, Cisticola exodus*. We recommend longer-term monitoring of these grassland birds in South Asian grasslands and farmlands in order to understand how they respond to ongoing land-use change as well as their relationship to mosaics of grassland-agriculture.

Our study also underscores the importance of historical data from natural history museums and archives for understanding the true magnitude of habitat loss on the local extirpation and decline of grassland birds. Many species in the tropics are being lost at unprecedented scales despite limited information about their natural history (Essl et al. 2026). Historical data from museums have been invaluable in advancing research and conservation for many species and ecosystems across the range but have so far been limited due to lack of access to data. To fill the vast knowledge gap on basic natural history of species in the Global South, we recommend utilizing historical datasets in tandem with modern surveys for the conservation and restoration of imperiled habitats like grasslands.

### Conservation Recommendations

Grasslands in South Asia are facing a multitude of threats owing to misclassification, conversion and lack of conservation prioritization (Ratnam et al. 2016b). Our study, integrating historical insights from collections and museums, shed valuable insights on the biodiversity collapse occurring in the Subansiri floodplain, and urge the immediate prioritization of conservation efforts in this landscape. In that regard, we offer the following recommendations-

1. **Map remaining wet grasslands**- while several efforts have been made to map the grasslands of India, much of them focus on the more open, semi-arid grasslands of central and western India (Madhusudan & Vanak 2022). A large reason for this is the fact that wet grasslands often exist alongside tropical moist deciduous forests and are also in the same climate envelope as forests. Thus, they are frequently mapped as forests. We urgently recommend a unified mapping effort that recognizes and maps all grassland types in India, including the wet grasslands of the Himalayan foothills.
2. **Scientific studies on long-term monitoring of birds**- In line with earlier recommendations (Rahmani et al. 2016), we reiterate the urgency in the long-term monitoring of grassland endemic species. This includes the genetic monitoring, population connectivity and species-habitat- disturbance relationships. With the shrinking of grassland species populations, there is a concern for lack of gene flow and increasing inbreeding within populations. Decline in genetic diversity can eventually lead to the local extinction of these species. However, there is no available genetic information for most of these grassland species (Reddy 2014). We recommend streamlined access to permits to map the extinction vulnerability and genetic diversity of grassland birds for urgent conservation management.
3. **Scientific assessment of grass and disturbance regimes**- Today, very little information is available about species-habitat-disturbance associations for most species of grassland birds of India, despite their threat status. Disturbance regimes such as fire, grazing and cutting are crucial for the persistence of grasslands and associated species because they create heterogeneity in the landscape (Fuhlendorf et al. 2006). However, how grassland birds of India respond to such disturbance regimes are as yet unknown (Rahmani 2016). We recommend an urgent need to study species- habitat relationships as well as species-disturbance regimes associations to incorporate this information into habitat management.

## ACKNOWLEDGEMENTS

We thank Anand Krishnan for helping collect data from Museum für Naturkunde Berlin and discussion on grassland birds, Dan Gordon for help with accessing Herbert Stevens specimens at the Newcastle Upon Tyne Museum, Mark Adams of NHM Tring for hosting SL at the museum, the UMN South Asia Library for access to historical archival materials, Rahul Khot and Vithoba for access to BNHS, VertNet for digital records of museum specimens at other NHMs. SL thanks Sakshi Nulkar for help with making figures, Vijay Ramesh for help with conducting archival research and sharing R codes for analyses. In the field, SL and JB thank members of Tesia Camp for logistical support. SL was funded through the Interdisciplinary Center for the study of Global Change (ICGC), Bell Museum of Natural History, Oriental Bird Club, American Ornithological Society. SR was funded by the U.S. National Science Foundation grant DEB-2321548. We recognize the complicated origins of natural history specimens and respectfully offer this Collections Acknowledgement: In aiming to document, preserve, and interpret biodiversity, we acknowledge the past histories and cultures from where the specimens originated. When using specimens to better understand the world’s biodiversity, we are obligated to uncover and document the entirety of the history of these collections. We are committed to creating an environment that welcomes and empowers people from historically underrepresented and marginalized communities to have equitable access to museum resources. Informed by this self-reflection and commitment, we continue our work to better understand the natural world, remember the past, and create a more equitable future.

## Supplementary Figures

**Fig S1:**
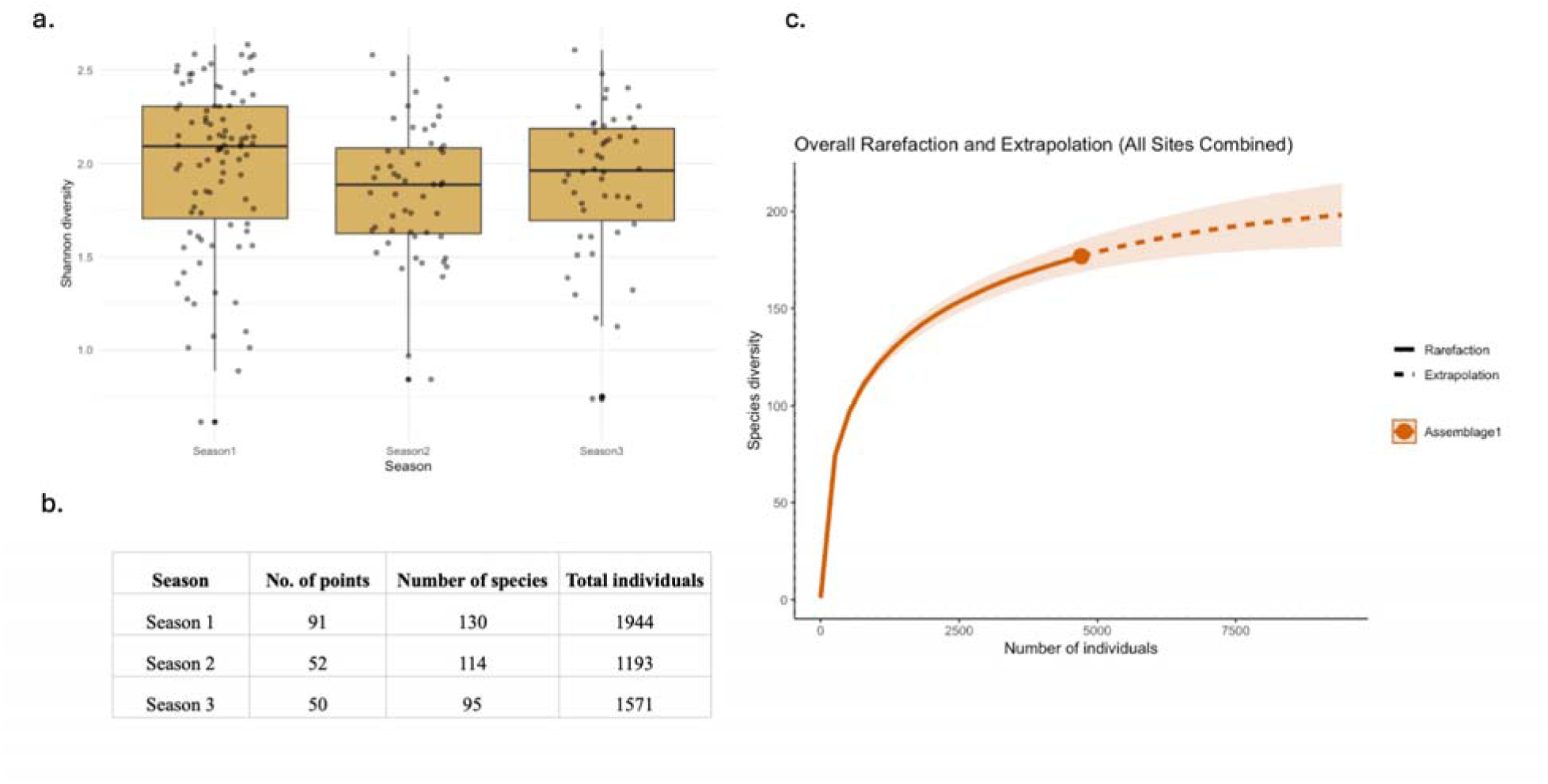
a and b. Shannon diversity and species richness were comparable across the three years of survey. C. Species accumulation curve with 99% completeness across all sites.

**Fig S2:**
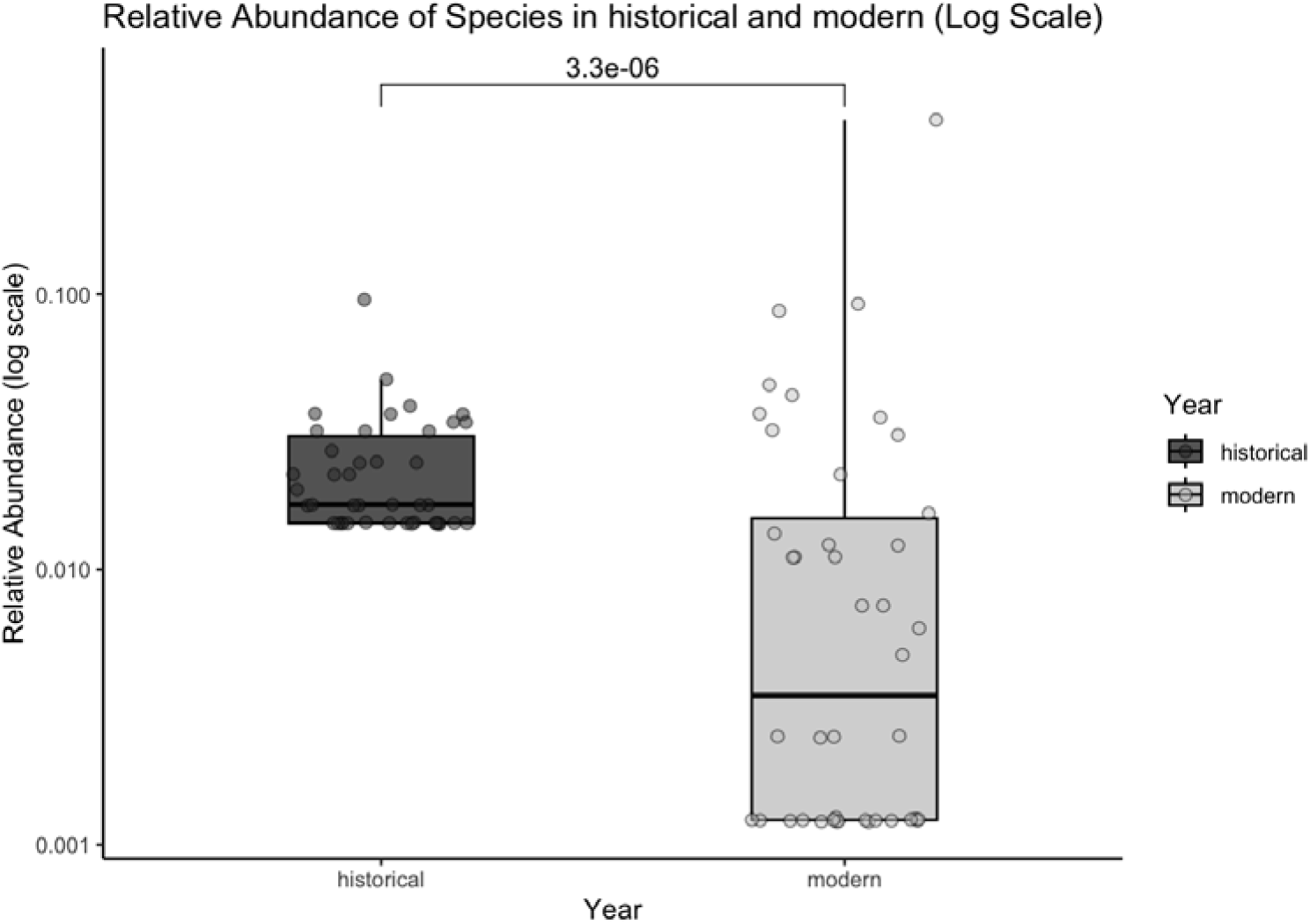
Relative abundance of species is significantly different, with reduction in abundance from historical to modern time period.

**Fig S3:**
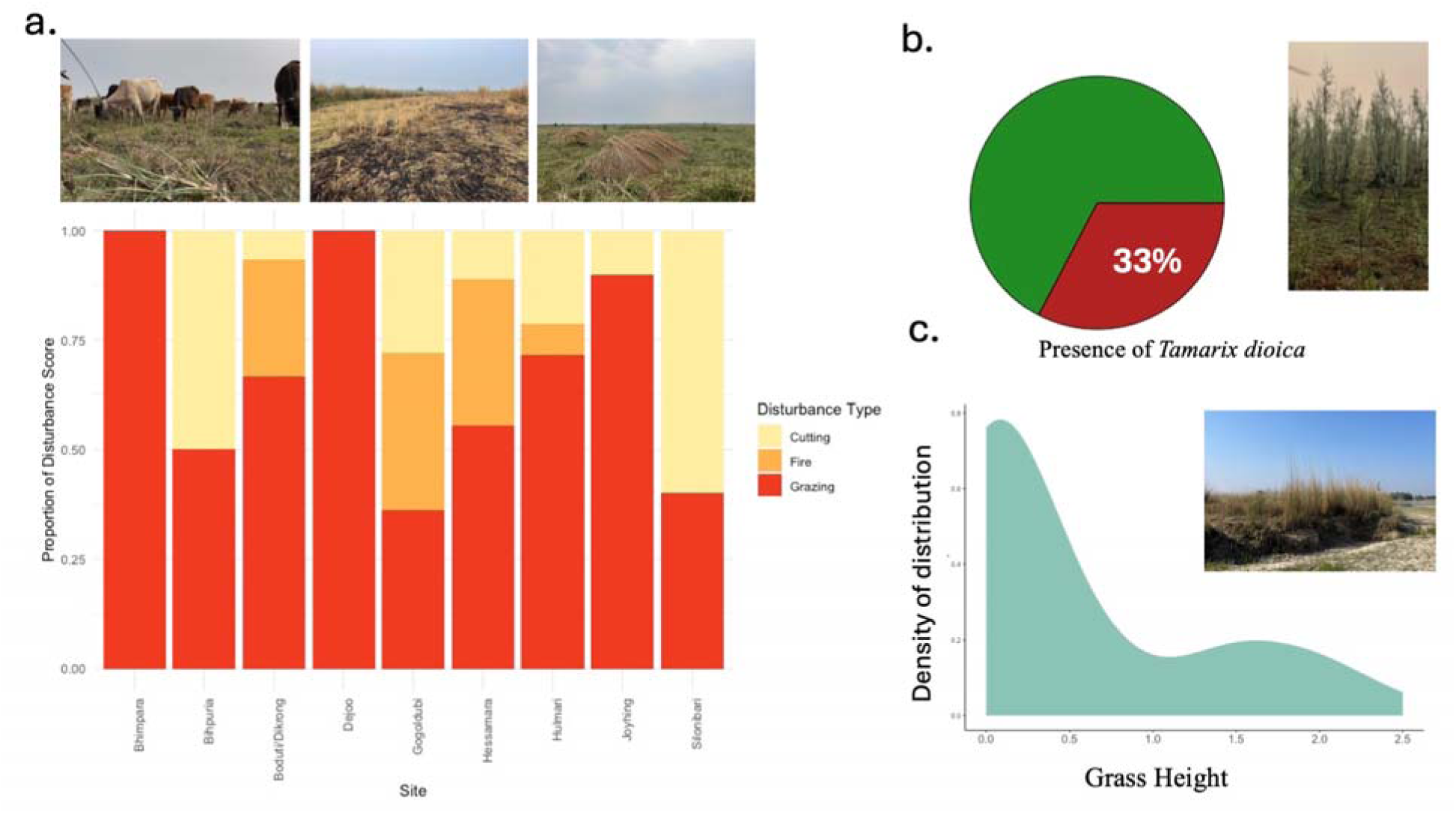
a. Proportion of disturbance regimes, such as grass cutting, fire and grazing across all our survey sites. 100% of our sites had evidence of grazing, 78% sites had evidence of grass cutting while 44% sites had evidence of fire. The picture panel above shows grazing, fire burning and grass cutting in our field sites. B. 33% of our sites had evidence of *Tamarix dioica.* C. Mean grass height was 0.55m.

